# Chemical Genetic Targeting of the LRRK2 GTPase Domain

**DOI:** 10.64898/2026.09.03.749217

**Authors:** Rachel E. Prorok, Lawrence Y. Zhu, Vickie Bowcut, Harry Wu, Keelan Z. Guiley, Johannes Morstein, Kevan M. Shokat

**Author notes:** Equal contribution (Rachel E. Prorok, Lawrence Y. Zhu).

## Abstract

Genetic variants throughout the multi-domain protein leucine-rich repeat kinase 2 (*LRRK2)* gene are the most common cause of autosomal dominant Parkinson’s disease, and the most prevalent Parkinson’s-associated LRRK2 variants enhance kinase activity. Hence, the kinase domain has been extensively targeted for therapeutic development. However, clinical progression of LRRK2 kinase inhibitors has been limited by on-target peripheral toxicities linked to strong kinase suppression, raising the question of whether targeting LRRK2 kinase activity through other means could offer a more tunable therapeutic window. LRRK2 is one of two proteins in the human proteome that possess a Roc-COR GTPase domain in tandem with a kinase domain, and the GTPase domain has been shown to intramolecularly regulate kinase output. Here, we ask whether the Roc-COR GTPase can be targeted as an alternative approach to downregulate kinase activity. We utilize a chemical genetic approach to sensitize LRRK2 to existing GTPase inhibitors and show that pharmacologically engaging the Roc-COR GTPase domain in cells decreases but does not fully inhibit LRRK2-mediated Rab10 T73 phosphorylation. This study lays the groundwork for future efforts directed at the development of pharmacological inhibitors for the LRRK2 GTPase, providing an alternative approach for therapies targeting LRRK2-driven Parkinson’s disease.

## Introduction

Mutations in Leucine-rich repeat kinase 2 (LRRK2) are the most common cause of autosomal dominant Parkinson’s disease (PD) (1–4) and recent studies have also demonstrated an association between non-coding LRRK2 variants and sporadic PD (5, 6). LRRK2 is a large, multidomain protein that possesses both a MAPKKK protein kinase and a Roc-COR (Ras of complex – C terminal of Roc) GTPase, in addition to several conserved protein-protein interaction domains (7, 8). Over 211 LRRK2 variants have been identified in PD patients, with the most frequent pathogenic mutations occurring in the GTPase (R1441C/G/H) and kinase (G2019S) domains. Almost all PD-associated LRRK2 variants cause a toxic gain-of-function in kinase activity, and increased LRRK2 kinase activity is also seen in idiopathic PD, underscoring the importance of this protein as a therapeutic target.

Although potent and selective LRRK2 kinase inhibitors have been developed, clinical translation of these agents have been hindered by preclinical safety findings demonstrating pulmonary and renal toxicities in non-human primates, a result of on-target LRRK2 inhibition outside the central nervous system (CNS) (9–11). Despite these challenges, the Phase 2b LUMA trial testing LRRK2 kinase inhibitor BIIB122 was completed but failed to achieve its primary endpoint of slowing disease progression in idiopathic PD patients. Excellent peripheral LRRK2 inhibition was observed as well as a modest reduction in the cerebrospinal fluid (CSF) biomarker phosphorylated Rab10 (pRab10), although the full data has not been published. While the ongoing Phase 2a BEACON trial will assess BIIB122 in LRRK2 variant carriers, it remains uncertain whether the doses required to achieve sufficient pRab10 inhibition in the CNS will be limited by peripheral consequences associated with LRRK2 kinase inhibition and LRRK2 knockout (12). The challenges of on target off-tissue toxicity by LRRK2 kinase inhibitors has prompted interest in exploring alternative therapeutic modalities, including LRRK2 degradation induced by proteolysis-targeting chimera (PROTAC) ARV-102, currently in Phase 1 clinical trials, and LRRK2 knockdown by the intrathecal delivery of antisense oligonucleotide (ASO) BIIB094, which was discontinued after completion of Phase 1 trials (13).

The strong genetic association between elevated LRRK2 kinase activity and PD risk supports efforts that directly target the kinase domain or reduce protein abundance. However, LRRK2 differs from canonical protein kinases in that its kinase activity is subject to multiple layers of autoregulation. Structurally, the cytosolic pool of LRRK2 exists in an autoinhibited conformation in which its N-terminal protein-binding domains dock across the C-terminal half of the protein, sterically occluding the catalytic cleft and maintaining the kinase lobes in an open, inactive conformation (14). Upon Rab-mediated membrane recruitment, the N-terminal domains undock to reveal the kinase catalytic cleft, and the C-terminus of the protein is permitted to rearrange such that the kinase lobes can adopt a closed, active conformation (15, 16) . In addition to this spatial regulation, genetic and biochemical evidence suggest the GTPase domain of LRRK2 also regulates kinase activity through a mechanism that remains incompletely understood (17, 18).

Mutations in the Roc-COR domain that decrease GTPase activity also enhance kinase activity, suggesting a relationship between nucleotide state of the Roc GTPase and kinase activation (16, 19–21). This regulatory paradigm is reminiscent of the relationship between a small GTPase and a downstream kinase, such as Ras and Raf, respectively, but for LRRK2 it is contained within a single polypeptide chain. Consistent with the small GTPase family, the Roc domain retains the conserved Ras-like GTPase fold, but it possesses relatively weak nucleotide affinity and its position between the N-terminal domains and the kinase domain constrains its conformational freedom. Moreover, the switch regions, which undergo a pronounced conformational rearrangement between GTP and GDP-bound states in small GTPases, are distinct in Roc: the Switch I region is longer and more flexible, and the Switch II region is buried at an interface with the neighboring COR-B domain (22, 23). Structural studies of the Roc-COR unit suggest that the GTPase domain may function as a non-canonical molecular switch that communicates activation state by propagating nucleotide-dependent conformational changes through surrounding LRRK2 domains, providing a potential mechanism through which the GTPase domain could allosterically regulate activation of the kinase domain (24, 25). We thus hypothesize that an inhibitor of the GTPase domain may serve as an underexplored alternative to direct kinase inhibition.

Until recently, GTPases have been considered “undruggable” due to their high nucleotide affinity and lack of allosteric pockets, but the discovery of the cryptic switch II pocket (SIIP) in the small GTPase K-Ras enabled the development and FDA-approval of three GTPase inhibitors, sotorasib, adagrasib, and daraxonrasib. These covalent inhibitors are dependent on the formation of a covalent bond with the oncogenic K-Ras(G12C) mutation and accordingly show no inhibition of the wild-type protein. The complexity of LRRK2s regulation, as well as the divergence of the Roc-COR unit from canonical small GTPases, make it difficult to predict how an inhibitor of the LRRK2 GTPase domain will impact kinase activity. Recently, we used chemical genetics to leverage the conservation of the SIIP in the Ras superfamily to target Ras-related GTPases with covalent K-Ras(G12C) SIIP inhibitors (26). We hypothesized that K-Ras(G12C) SIIP inhibitors could similarly be used to target the Roc-COR GTPase in LRRK2 as a model to understand whether binding this domain in LRRK2 would be a viable alternative strategy to existing kinase domain-targeting drug candidates. We utilized a chemical genetic approach to engineer a K-Ras(G12C)-equivalent mutation in the LRRK2 Roc domain, T1343C, and demonstrated binding by known K-Ras(G12C) inhibitors. Building upon this established framework, we introduced multiple mutations contained within the SIIP to replicate key drug stabilizing interactions and observed enhanced binding efficiency. The resulting LRRK2 construct possesses a neo-SIIP that enables chemical genetic targeting of LRRK2 by the clinical K-Ras inhibitor divarasib *in vitro* and in cells. Here, we demonstrate for the first time that targeting the LRRK2 GTPase domain achieves effective target inhibition, with distinct effects on LRRK2 conformation and pRab10 compared to established kinase inhibitors.

## Results

### Mutation of the LRRK2 GTPase protein is required for engagement by K-Ras(G12C) inhibitor

Sequence identity of the Roc domain with K-Ras4B is low (21.9%) and the residues that shape the SIIP are not conserved in LRRK2 (Figure 1B), so we first investigated whether the SIIP of Roc could be targeted despite the lack of high sequence homology. We tested three different SIIP ligands for covalent engagement with Roc_ext_(T1343C), a construct of the Roc domain engineered with a single non-native cysteine at residue T1343 to mimic the somatic mutation in K-Ras(G12C) (27). Unless stated otherwise, all Roc_ext_ constructs used in this study lacked the single native cysteine (C1465S) to facilitate uniform labeling. Using whole protein mass measurements, we found that Roc_ext_(T1343C) exhibited the expected mass shift when incubated in the presence of sotorasib, divarasib, and adagrasib, demonstrating that the T1343C mutation in LRRK2 is available for covalent attack of the acrylamide warhead (Figure 1D, E). Reversion of T1343C to the native threonine ablated covalent labeling, confirming the specificity of the modification. However, Roc_ext_(T1343C) was only partially labeled after 120 minutes, suggesting the lack of conservation of key residues around the SIIP prohibits effective ligand binding.

**Figure 1.**
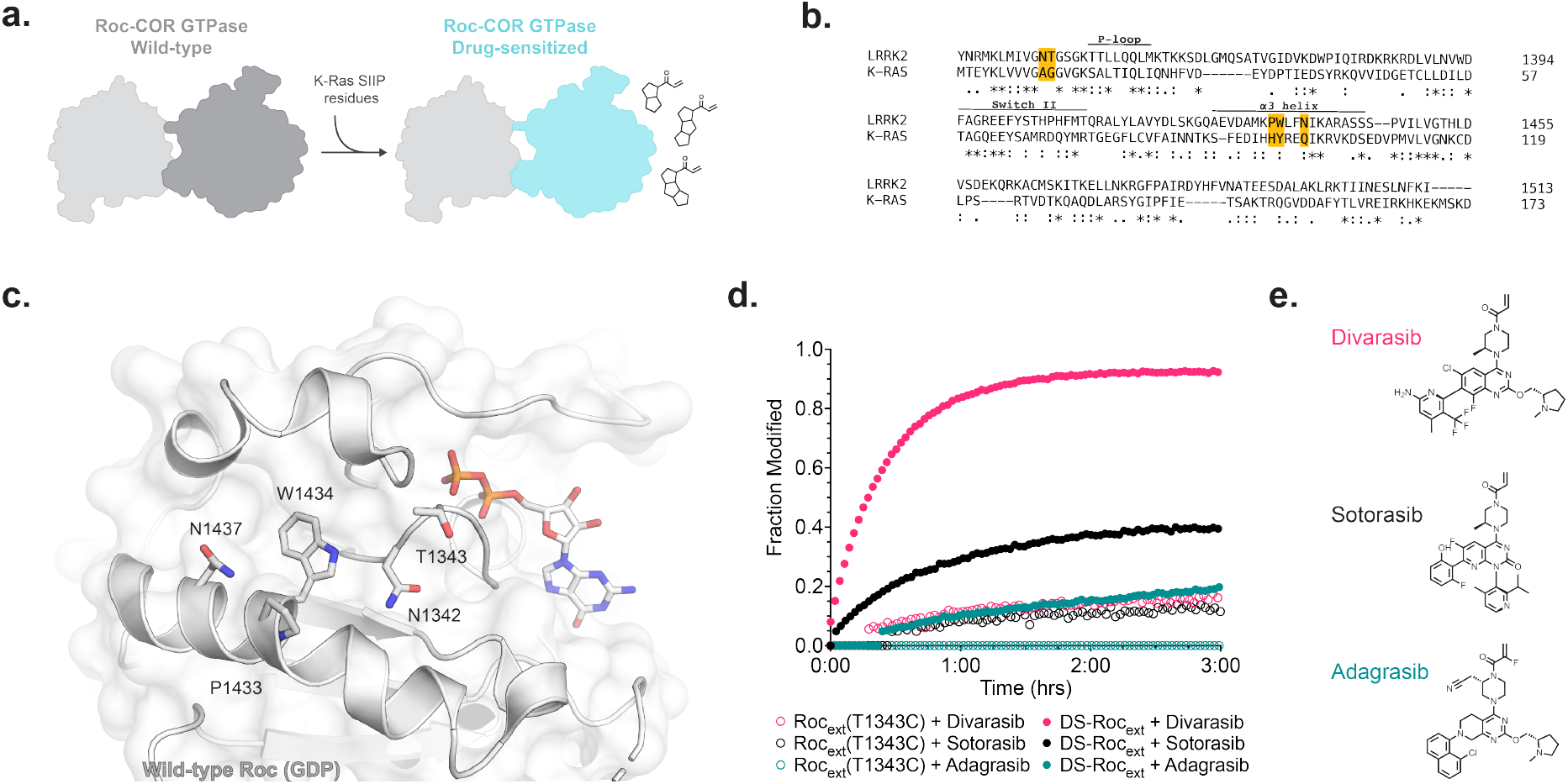
Drug-sensitizing Roc to K-Ras(G12C) inhibitors requires five Switch II pocket mutations. **A)** Schematic of drug-sensitizing the Roc-COR GTPase. **B)** Sequence alignment of LRRK2 and K-Ras. Switch II pocket residues mediating ligand:protein interactions between sotorasib and K-Ras are highlighted in yellow. **C)** CryoEM structure of GDP-bound Roc highlighting key Switch II pocket side chains (PDB: 7LHW). **D)** Time-dependent covalent labeling of Roc_ext_ and DS-Roc_ext_ by divarasib, sotorasib, and adagrasib. **E)** Chemical structures of K-Ras(G12C) inhibitors tested against Roc_ext_.

To improve labeling of Roc_ext_(T1343C), we next sought to more closely recapitulate the non-covalent stabilizing interactions between K-Ras(G12C) and SIIP ligands. In the co-crystal structure of sotorasib and K-Ras(G12C) (PDB:6OIM), the residues H95, Y96, and Q99 in the α3 helix are key to forming and stabilizing sotorasib within the cryptic pocket (Figure 1C). In addition, we hypothesized that labeling kinetics could be further improved by removing steric bulk surrounding T1343C, increasing accessibility of the nucleophile. As such, we introduced a total of four mutations - N1342A, P1433H, W1434Y, and N1437Q - into Roc_ext_(T1343C) to form a drug-sensitized construct of Roc, DS-Roc_ext_. Importantly, P-loop residues required for nucleotide binding and switch II helix residues required for GTP hydrolysis remained unmodified (28). We then used whole protein mass measurements to assess labeling of DS-Roc_ext_ with divarasib, sotorasib, and adagrasib (29–31). We found that DS-Roc_ext_ underwent nearly complete labeling with divarasib and sotorasib within 120 minutes, validating the approach of leveraging non-covalent protein-ligand interactions to improve binding (Figure 1D). To our surprise, we found that adagrasib labeled relatively poorly, despite a key prerequisite histidine P1433H (H95 in K-Ras) having been introduced (32–34). As a result, we focused on divarasib for all subsequent experiments due to its superior labeling relative to adagrasib and sotorasib.

We assessed the relative contribution of each individual mutation in DS-Roc_ext_ to ligand binding. Reversion of any one of the three mutations on the α3 helix (H1433P, Y1434W, Q1437N) significantly reduced labeling by divarasib, whereas reversion of the P-loop mutation N1342A had no effect on maximum labeling but slowed labeling kinetics (Supplementary Figure 1). Taken together, these data indicate that non-covalent interactions between the ligand core and SIIP residues significantly contribute to ligand binding, whereas p-loop residues likely influence reactivity of the electrophilic warhead.

### DS-Roc_ext_ forms a complex resembling K-Ras(G12C) with SIIP inhibitors

To evaluate whether the introduced mutations in DS-Roc_ext_ recreate the divarasib binding pose observed in K-Ras(G12C), we solved an x-ray co-crystal structure of the DS-Roc_ext_:divarasib complex with an overall resolution of 2.3 Å (PDB: 9C76) (Figure 2A). The protein-drug complex crystallized as a domain-swapped homodimer, with the β1, α1, β2, β3, and α2 elements from one monomer associating with the β4, α3, β5, α4, β6, and α5 elements of another. This is consistent with previously reported apo structures of Roc_ext_ (rmsd 3.47 Å) (PDB: 6XAF) (27). Although the biological relevance of this arrangement has been debated in the literature (35), it does not preclude our analysis of the binding pose of divarasib as it primarily forms contacts with the α3 helix.

**Figure 2.**
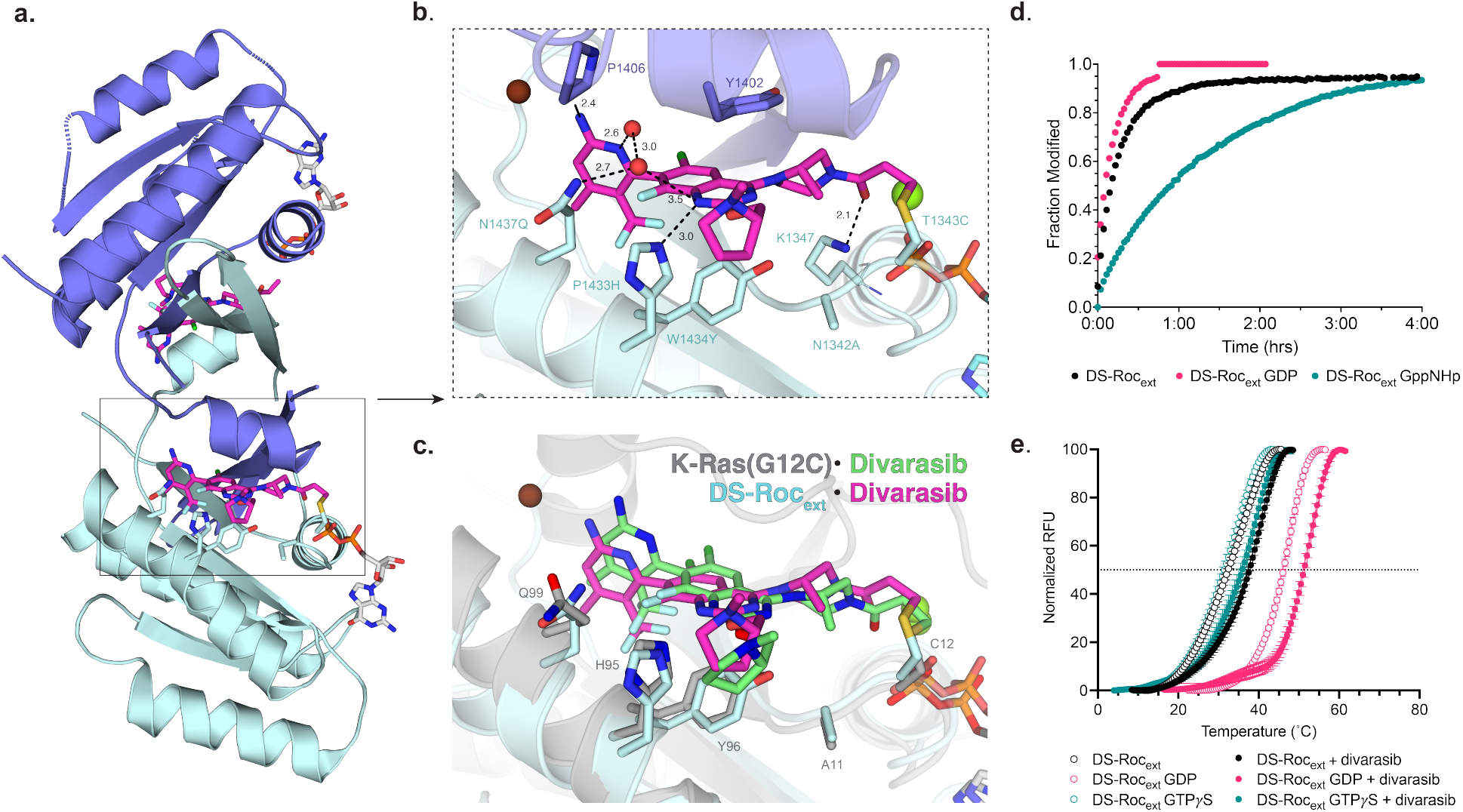
Divarasib binds DS-Roc_ext_ in the Switch II pocket and stabilizes GDP. **A)** X-Ray crystal structure of the chain-swapped DS-Roc_ext_ homodimer and divarasib (PDB: 9C76). **B)** Inset of divarasib in the Switch II pocket. Distances are in Å. **C)** Overlay of co-crystal structures of DS-Roc_ext_·Divarasib and K-Ras(G12C)·Divarasib (PDB: 9DMM). **D)** Time-dependent covalent labeling of Roc_ext_ and DS-Roc_ext_ by divarasib in the presence of saturating concentrations of nucleotide. **E)** Differential scanning fluorometry of DS-Roc_ext_·Divarasib adduct. Refer to Supplementary Table 2 for Tm.

Close examination of divarasib reveals several ligand-protein interactions that replicate what is observed in K-Ras(G12C):divarasib. The histidine side chain of introduced residue P1433H (K-Ras H95) adopts the same “in” conformation observed in the K-Ras(G12C):divarasib complex, positioning it to participate in hydrogen bonding with the quinazoline core of divarasib at a distance of 3.0Å (Figure 2B). Since adagrasib also relies on this His-dependent interaction, it is notable that sotorasib, which binds with the histidine side chain oriented toward solvent and lacks His-mediated hydrogen bonding, demonstrates more efficient protein labeling than adagrasib (PDB:6OIM) (36). Located at the base of the pocket, the substitution W1434Y (K-Ras Y96) adopts a similar pose to the equivalent residue in K-Ras(G12C):divarasib. The introduced residue N1437Q (K-Ras Q99) does not form obvious interactions with divarasib but instead coordinates a water molecule that participates in a hydrogen bonding network with the pyridine 5-NH_2_ and the peptide backbone of the switch II (SII) loop. This differs from the observed pose of Q99 in K-Ras(G12C):divarasib, where the amide nitrogen points away from divarasib and exhibits no hydrogen bonding to the drug (Figure 2C).

In addition to the introduced residues, several native SIIP residues interact with divarasib. The tri-fluoro group of divarasib resides deep within a hydrophobic pocket composed by V1340, I1438, Y1415 (Supplementary Figure 2A). In the switch II loop, the side chain of Y1402 participates in pi-pi stacking with the quinazoline core. At the back of the pocket, K1347 (K-Ras K16) forms a hydrogen bond with the acrylamide carbonyl at a distance of 2.1Å in an interaction mirroring what is observed in the K-Ras(G12C):divarasib complex (Figure 2B). The presence of multiple interactions between the ligand and both native and non-native protein residues, in conjunction with biochemical evidence from labeling of Roc_ext_(T1343C), highlights the importance of secondary interactions in facilitating reversible affinity in the SIIP of DS-Roc_ext_.

### SIIP engagement modulates nucleotide state preference of DS-Roc_ext_

Most SIIP inhibitors, including sotorasib and adagrasib, exhibit selective covalent labeling of GDP-bound K-Ras(G12C) with negligible labeling of the GTP-bound state (32, 36–38). To evaluate whether divarasib preferentially labels DS-Roc_ext_ in a specific nucleotide state, we compared covalent labeling under GDP or GppNHp-saturated conditions. As observed with K-Ras(G12C), divarasib preferentially reacts with DS-Roc_ext_ under GDP-saturated conditions, however in contrast to K-Ras(G12C), DS-Roc_ext_ labeling still occurs at a reduced but measurable rate in GppNHp-saturated conditions (Figure 2D). We speculate that the discrepancy in nucleotide preference is due to a difference in how nucleotide state is translated to the conformation of switch II in the Roc GTPase compared to K-Ras. Unlike K-Ras, where the switch II undergoes a pronounced disorder-to-order transition upon GppNHp binding, structural alignment of the Roc domain in GDP and GppNHp-bound LRRK2 shows the switch II remaining relatively ordered in both states (39).

To study the effect of divarasib binding on DS-Roc_ext_ stability, we performed differential scanning fluorimetry (DSF) in nucleotide-free, GDP-saturating, or GTPγS-saturating conditions. We found that divarasib significantly increases the thermal stability of DS-Roc_ext_ by an average of 5.3◦C for all nucleotide conditions tested (Figure 2E), indicating that divarasib does not preferentially stabilize a specific DS-Roc_ext_ nucleotide state. These findings, in combination with nucleotide dependence LCMS-based studies, demonstrate that divarasib may not possess a strong nucleotide preference for binding the SIIP in DS-Roc_ext_, diverging from the well-characterized binding mode of first generation SIIP inhibitors to K-Ras(G12C) (38, 40). The ability to engage both GDP and GTP states of K-Ras (G12C) has been recently reported for second generation inhibitors (41).

### LRRK2-DS displays elevated kinase activity in cells

Having demonstrated both biochemical and structural evidence for sensitivity of DS-Roc_ext_ to divarasib, we subsequently investigated whether the same set of mutations could be used to sensitize full-length LRRK2 to SIIP ligands. To first evaluate the impact of adding five drug-sensitizing mutations into LRRK2, we utilized a HEK293 cell overexpression system to transiently express full-length drug-sensitized LRRK2 (LRRK2-DS) or wild-type LRRK2 (LRRK2-WT) isogenic controls. We assessed LRRK2 kinase activity by measuring the biomarker phosphorylation site LRRK2 S935 and the downstream LRRK2-dependent phosphorylation site Rab10 T73 by western blot (42, 43). When compared to LRRK2-WT, we observed a significant decrease in LRRK2 pS935 signal and an increase in Rab10 pT73 signal for LRRK2-DS, indicating that introduction of the drug-sensitizing mutations increases LRRK2 kinase activity (Figure 3A-C).

**Figure 3.**
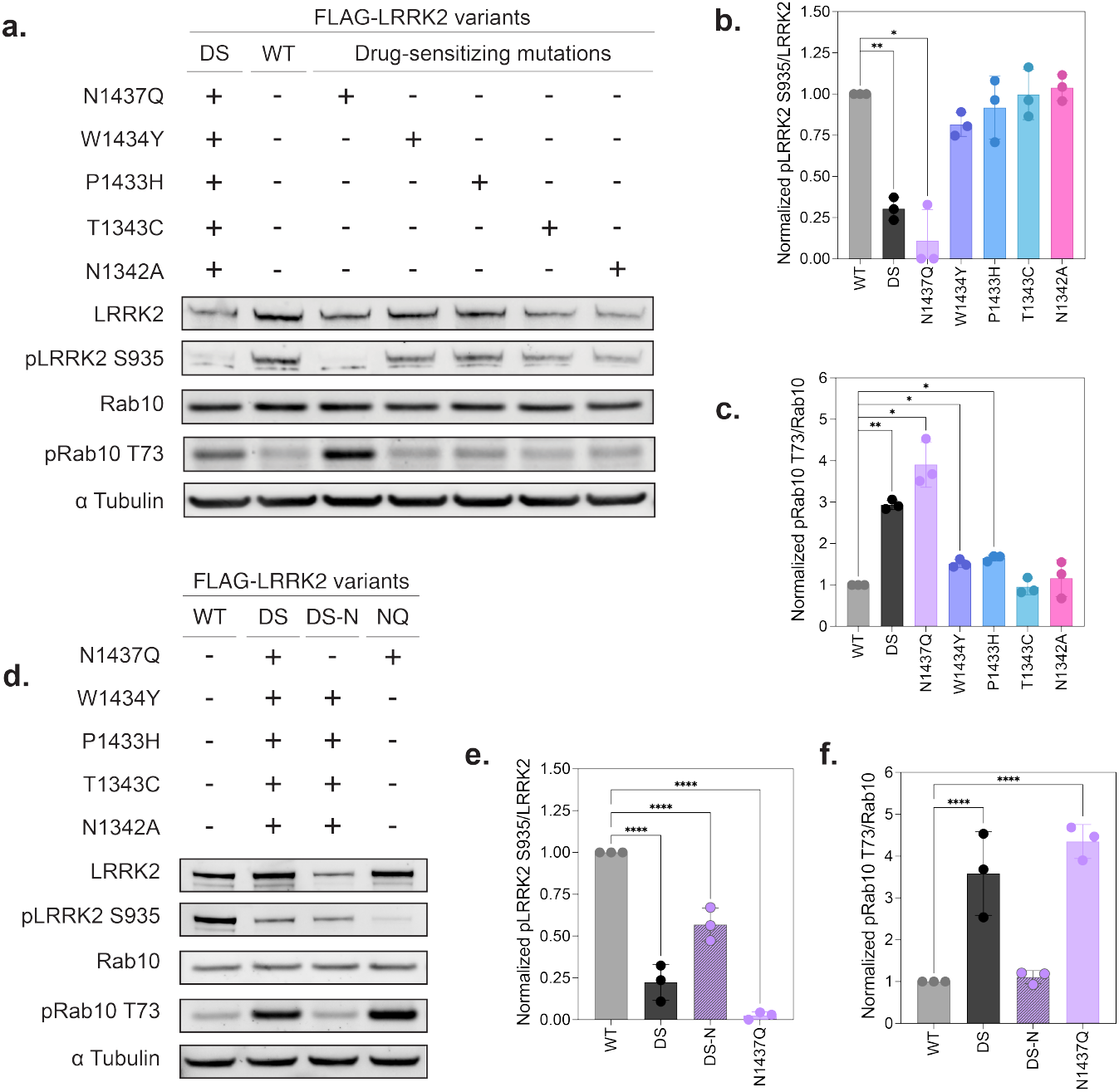
Drug-sensitizing mutation N1437Q increases kinase activity of DS-LRRK2. **A)** Immunoblot analysis of HEK293T cells overexpressing FLAG-LRRK2 containing drug-sensitizing mutation variants. **B)** Immunoblot quantification of LRRK2 S935 phosphorylation for immunoblot shown in Figure 3A. **C)** Immunoblot quantification of Rab10 T73 phosphorylation for immunoblot shown in Figure 3A. **D)** Immunoblot analysis of HEK293T cells overexpressing wild-type (WT), drug-sensitized (DS), drug-sensitized with native N1437 (DS-N), or N1437Q (NQ) FLAG-LRRK2. **E)** Immunoblot quantification of LRRK2 S935 phosphorylation for immunoblot shown in Figure 3D. **F)** Immunoblot quantification of Rab10 T73 phosphorylation for immunoblot shown in Figure 3D. Immunoblot analyses (B-C, E-F) are mean +/-SD (n=3 independent experiments).

Prior to investigating the impact of divarasib treatment of LRRK2-DS, we first investigated the unexpected activation of LRRK2-DS by the mutations introduced to sensitize to divarasib. The LRRK2-DS phosphorylation pattern resembles previously reported LRRK2 pS935 and Rab10 pT73 signals from PD-associated LRRK2 variant LRRK2-R1441C (19). Located at the terminus of the α3 helix in the Roc GTPase, the R1441C variant is hypothesized to activate LRRK2 by destabilizing the Roc:COR-B interface, causing LRRK2 to undergo a conformational shift away from its autoinhibited conformation and promoting activation of the kinase domain (16). In contrast, MD simulations suggest the widely prevalent kinase domain variant G2019S reinforces the closed, active conformation of the kinase domain by stabilizing the DYG motif (14, 44). Given that three of the five mutations used to generate LRRK2-DS are located in the α3 helix, we considered whether the drug-sensitizing mutations activate the kinase domain through a similar mechanism as R1441C. Structural analysis suggests R1441 forms contacts with residues in COR-B that are not permissible after LRRK2 adopts its activated conformation (16) Applying this framework to the drug-sensitizing mutations, we utilized the cryo-EM structure of full-length, inactive LRRK2 (PDB 7LHW) to identify all residues in COR-B that are located within 5Å of the drug-sensitizing mutation positions in Roc. While no COR-B contacts were identified for residues T1343 or N1342, we found that P1433, W1434, and N1437 are positioned to contact several residues in COR-B (**S**upplementary Figure 3). We became specifically interested in the contacts formed by residue N1437 because of its proximity to a COR-B loop, and because N1437D/H have been characterized as highly penetrant PD-associated variants. We asked whether the drug-sensitizing mutation N1437Q mimics the kinase-activating effect of these PD-associated variants and found that N1437Q produces the same LRRK2 S935 and Rab10 T73 phosphorylation state as N1437H (Supplementary Figure 4).

To complement our structural analysis and test whether N1437Q is the primary residue responsible for LRRK2-DS activation, we transfected HEK293 cells with LRRK2 containing each of the individual drug-sensitizing mutations and assessed kinase activity (Figure 3A-C). Only N1437Q mirrors the effect of LRRK2-DS in the absence of additional mutations. Accordingly, reversion of glutamine to the native asparagine in LRRK2-DS-N partially recovers LRRK2 pS935 and fully corrects the increase in Rab10 pT73, further corroborating N1437Q as the activator of elevated kinase activity in LRRK2-DS (Figure 3D-F). Supported by previous structural analyses of active and inactive LRRK2, this data emphasizes a critical role for residue N1437 in stabilizing the Roc:COR-B interdomain interface and suggests that mutations disrupting the interface can activate the kinase domain (15).

### Switch II occupancy by divarasib alters LRRK2 cellular activity

We next turned to interrogate the impact of pharmacologically engaging the Switch II pocket, which is located at the interface of the Roc and COR-B, of LRRK2-DS with divarasib. To assess this, we transiently overexpressed LRRK2-DS in HEK293 and measured phosphorylation of LRRK2 S935 and Rab10 T73 following treatment with divarasib. Strikingly, divarasib treatment decreases both Rab10 pT73 and LRRK2 pS935 signal compared to vehicle-treated LRRK2-DS (Figure 4A-C). Compared to the type I LRRK2 kinase inhibitor MLi-2, divarasib similarly reduces LRRK2 pS935 but only partially reduces Rab10 pT73 to a level found in LRRK2-WT expression conditions, whereas MLi-2 completely ablates Rab10 phosphorylation (Figure 4A-C) (45, 46). The phosphorylation pattern of S935 and T73 produced by SIIP occupancy is not explained by either established mode of LRRK2 kinase inhibition.

**Figure 4.**
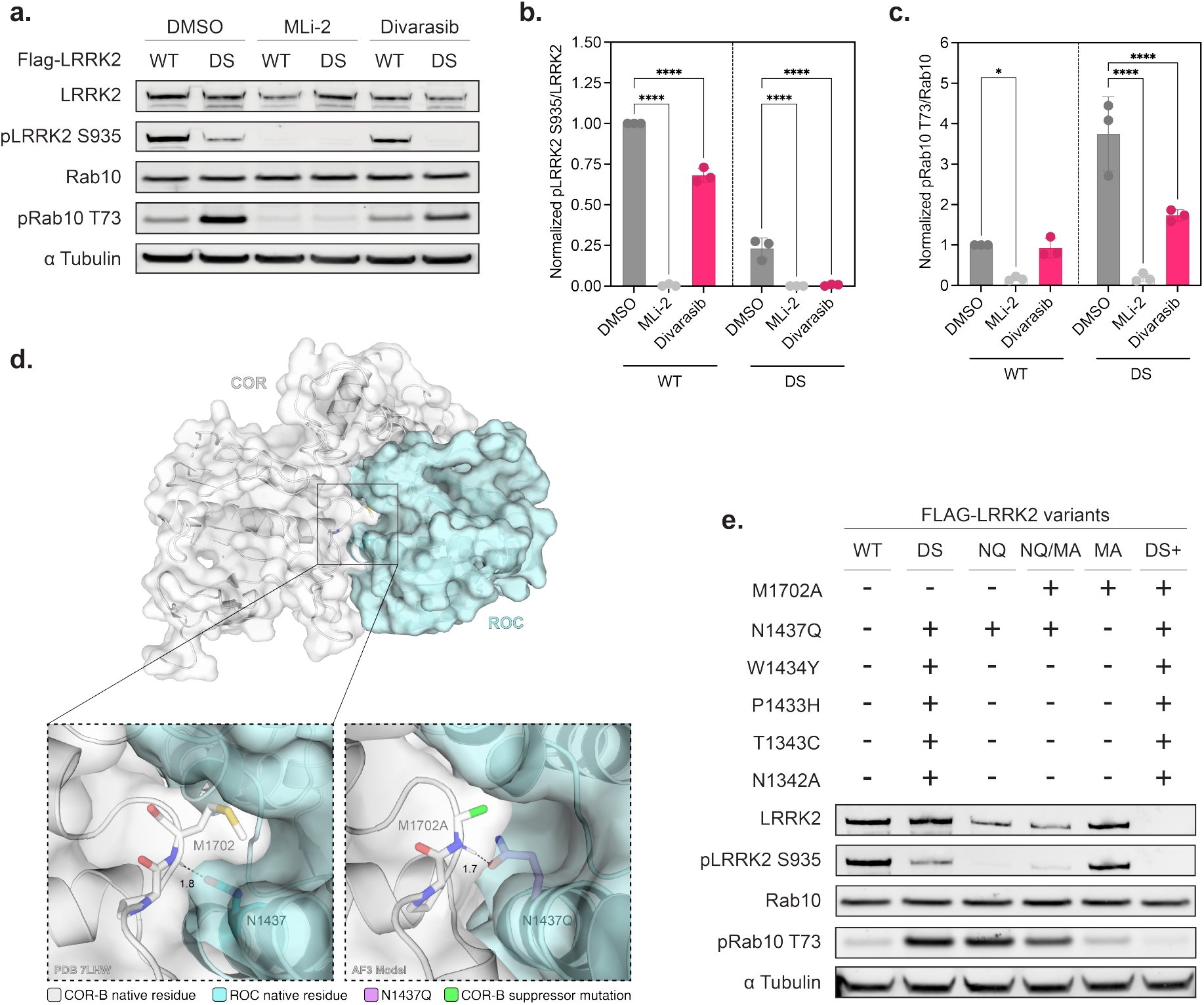
Pharmacological and genetic intervention at the Roc:COR interface restrains LRRK catalysis. **A)** Immunoblot analysis of HEK293T cells overexpressing wild-type (WT) or drug-sensitized (DS) FLAG-LRRK2 following 8 hour treatment with 10 µM Divarasib or 90 minute treatment with 100 nM MLi-2. **B)** Immunoblot quantification of LRRK2 S935 phosphorylation for immunoblot shown in Figure 4A. **C)** Immunoblot quantification of Rab10 T73 phosphorylation for immunoblot shown in Figure 4A. **D)** Surface representation of the Roc-COR GTPase with inset of interactions between N1437 and M1702. Distances are in Å (WT PDB: 7LHW, N1437Q/M1702A AF3 model). **E)** Immunoblot analysis of HEK293T cells overexpressing wild-type (WT), drug-sensitized (DS), N1437Q (NQ), N1437Q/M1702A (NQ/MA), M1702A (MA) or drug-sensitized with M1702A (DS+) FLAG-LRRK2 variants. Refer to Supplementary Figure 5b for immunoblot quantification. Immunoblot quantifications (B-C) are mean +/-SD (n=3 independent experiments).

Type I inhibitors, which stabilize a closed, DYG-in kinase domain and a correspondingly non-autoinhibited global conformation, are associated with pS935 dephosphorylation (47). Type II inhibitors, on the contrary, bind the kinase domain in an open, DYG-out conformation and maintain LRRK2 in a global autoinhibited state, preserving S935 phosphorylation (48, 49). Both models reduce Rab10 pT73 from active-site occlusion. Divarasib treatment phenocopies the reduction of pS935 associated with Type I inhibition, but it also reduces Rab10 pT73, which is incompatible with the Type I model in which the kinase domain is closed and active. Alternatively, if divarasib followed the Type II model and stabilized an autoinhibited conformation with the kinase in an open, inactive conformation with decreased Rab10 pT73 phosphorylation, then pS935 would be preserved. To dissect whether the residual Rab10 pT73 reflects active-site-dependent LRRK2 kinase activity, we performed a sequential treatment of divarasib followed by MLi-2 and found that MLi-2 was indeed able to eliminate the remaining Rab10 pT73 signal (Supplementary Figure 5). Together, these data suggest that chemical genetic engagement of the Roc SIIP dissociates the global conformational state reported by S935 from its cognate catalytic output, suggesting the GTPase module regulates kinase activity through multiple mechanisms.

### COR-B second-site mutation attenuates LRRK2-DS kinase activity

The SIIP is located at the Roc:COR-B interface, which we hypothesize is destabilized in LRRK2-DS by N1437Q. Since this interface couples the Roc GTPase domain to the kinase domain, we reasoned that introduction of an interface-restabilizing mutation in COR-B may compensate for N1437Q-driven kinase activation and produce a similar effect as divarasib binding. To test this, we first identified COR-B residues that contact N1437 (13). Positions M1702 and P1701 in COR-B are located within 4Å of N1437, and the oxygen of N1437 forms a hydrogen bond with the backbone amide of M1702 at distance of 1.8Å (Figure 4D). Mutation of asparagine to glutamine could disrupt this interaction by increasing the length and flexibility of the side chain, leading to suboptimal positioning of the glutamine carbonyl for hydrogen bonding. Additionally, the added length of glutamine may sterically perturb packing at the Roc:COR-B interface, augmenting instability at the interface and shifting equilibrium to further favor the activated state of LRRK2.

To explore these hypotheses, we reduced the steric bulk of the COR-B side chain by introducing M1702A as a second-site suppressor mutation with N1437Q, and we subsequently assessed LRRK2 kinase activity in cells (Figure 4D, E). On its own, M1702A has no significant effect on LRRK2 pS935 or Rab10 pT73 levels. In combination with N1437Q, M1702A decreases Rab10 pT73 levels by 39.6% relative to N1437Q, replicating the catalysis-attenuating effect we observed in our chemical genetic approach. This result is consistent with our expectation that poor interface packing may contribute to the kinase-activating effect of drug-sensitizing mutation N1437Q. The similarities of the divarasib and second-site suppressor phenotypes indicate that the Roc:COR-B interface may play a pivotal role in governing both global conformational state of LRRK2 and its kinase catalytic output – two mechanisms through which the GTPase module can regulate kinase activation.

## Discussion

LRRK2 is a critical therapeutic disease target for Parkinson’s disease as it represents the most common familial cause of PD. Multiple clinical trials of LRRK2 kinase inhibitors in idiopathic patients have been disappointing, with the Phase 2a BEACON trial of the LRRK2 kinase inhibitor BIIB122 in LRRK2 variant carriers set to read out in early 2027. Preclinical studies of LRRK2 kinase inhibitors have identified on-target off-tissue toxicity as a liability which could limit dosing in ongoing trials. As a potential alternative to current strategies in the clinic, we investigated whether a different domain in LRRK2, the Roc-COR GTPase could be an alternative approach to inhibition of LRRK2 kinase function. The crosstalk between its two catalytic domains, the Roc-COR GTPase and the kinase, remains underexplored in cells.

Here, we utilized orthogonal approaches to demonstrate the regulatory relationship between the Roc-COR GTPase and activity of the kinase domain. First, we leveraged the presence of the cryptic SIIP in the Roc domain to generate a chemical genetic tool that enabled targeting of the GTPase with existing chemical matter. We showed precise recapitulation of the K-Ras(G12C) SIIP and divarasib binding pose in a solved crystal structure of the DS-Roc_ext_:divarasib complex. Following additional biochemical and cellular characterization of the chemical genetic tool LRRK2-DS, we demonstrated that pharmacological engagement of the Roc SIIP alters kinase activity, supporting the Roc GTPase as an allosteric regulator of kinase activity. In parallel, we performed a structural analysis of our drug-sensitized construct LRRK2-DS and identified a compensatory mutation in COR-B that attenuates the kinase-activating effect of drug-sensitizing mutation N1437Q, reproducing the kinase-attenuating effect observed with divarasib.

The LRRK2 S935 and Rab10 T73 phosphorylation signature resulting from occupancy of the SIIP is distinct from the phosphorylation patterns associated with either type I or type II kinase active-site inhibition. Treating DS-LRRK2 overexpressing cells with divarasib causes a paradoxical decrease in LRRK2 pS935 with a decrease in pRab10, which has previously been well characterized as a measure of LRRK2 catalytic output (43, 50, 51). On the other hand, pS935 is not a direct measure of catalytic output but rather a marker of global LRRK2 conformation, with preservation of pS935 occurring when LRRK2 is in an autoinhibited state (52). Phosphorylation of S935 recruits LRRK2 to the molecular scaffold protein 14-3-3, which stabilizes LRRK2 in an autoinhibited conformation in the cytoplasm (53, 54). A recent cryo-EM structure of the LRRK2:14-3-3 complex shows that a 14-3-3 dimer engages LRRK2 through two interaction sites: the phosphorylated S910/S935 loop and the COR-A and COR-B domains (55). Tool mutations that disrupt the COR-A and COR-B interaction interfaces, as well PD-associated variants that disrupt the autoinhibited conformation, cause 14-3-3 dissociation and expose the S910/S935 to phosphatases (55–57).

Based on this model, we hypothesize that divarasib binding disrupts the autoinhibited conformation by causing a conformational shift in the GTPase module, breaking the COR:14-3-3 interactions and promoting pS935 dephosphorylation (16). However, the attenuation of pRab10 signal upon divarasib treatment suggests that there is a separate mechanism through which the Roc-COR domains constrain kinase activity beyond their control of the global autoinhibited conformation. We cannot yet distinguish whether this constraint is imposed by conformational restriction of the kinase domain or an effect on substrate engagement, though a cryo-EM structure of LRRK2-DS bound to divarasib may help resolve this.

Both our pharmacological and genetic approaches to targeting the GTPase domain localize regulation of the kinase domain to the Roc:COR-B interface. Both approaches resulted in attenuation of N1437Q-driven kinase hyperactivation by 40-50% without a corresponding increase in pS935 levels, demonstrating uncoupling of global conformational state from catalytic output. Additionally, our identification of the second-site suppressor mutation M1702A supports a model in which our drug-sensitizing mutation N1437Q as well as the PD-associated variants N1437D/H activate LRRK2 by destabilizing the interface and releasing GTPase module constraints on kinase activity, mirroring the mechanism of PD variant R1441C (16). By identifying a set of complementary mutations as well as a chemical genetic model for drug binding at the Roc:COR-B interface that both move Rab10 pT73 levels toward those observed in wild-type cells, rather than full inhibition, we demonstrate that the Roc-COR GTPase acts as a tunable regulator of kinase activation rather than a binary switch. An instructive precedent for restraining kinase activity as an approach distinct from complete kinase inhibition comes from mTORC1, where allosteric regulation of mTORC1 by rapamycin only blocks 50% of substrate 4E-BP1 phosphorylation while completely inhibiting phosphorylation of S6K (58, 59). Much like LRRK2, the serine/threonine kinase domain of mTORC1 is embedded within extensive regulatory domain architecture that enables catalytic output to be tightly controlled by the integration of incoming nutrient and hormonal signals (60). We hypothesize that the regulatory architecture that governs mTORC1 activity and enables partial catalytic restraint by rapamycin engagement may similarly enable LRRK2 to be only partially restrained by engagement of the Roc-COR GTPase. Our results highlight the feasibility of targeting the Roc-COR GTPase via the SIIP to inhibit LRRK2 pathological activity while preserving basal Rab phosphorylation. Collectively, these advances warrant further investigation into the therapeutic potential of targeting the Roc-COR GTPase as an alternative to direct kinase inhibition.

## Methods

### Recombinant protein expression and purification

Roc GTPase constructs were constructed utilizing codon optimized DNA fragments synthesized by Twist Biosciences, and inserted into the pJExpress411 vector utilizing Gibson assembly. These constructs were expressed as described previously (20). Briefly, N-terminal-hexahistidine tagged Roc_EXT_ constructs were transformed into BL21 cells (NEB) and grown to 0.6 OD_600nm_ prior to induction with 5mM IPTG. Induction continued overnight at 20 °C. Then, bacterial pellets were isolated with centrifugation and lysed using wash buffer (30 mM HEPES (pH 7.4), 500 mM NaCl, 10 mM MgCl_2_, 20 mM imidazole, 50 μM GDP, and 10% (vol/vol) glycerol) supplemented with 1% v/v PMSF. The bacterial suspension was lysed using sonication and subjected to Ni-NTA resin (Thermo Fisher Scientific) batch chromatography. No elution with imidazole was performed, but instead the resin was washed with TEV buffer (30 mM HEPES (pH 7.4), 250 mM NaCl, 2 mM MgCl_2_, 5 mM BME, 100 uM GDP, and 10% glycerol), followed by addition of His-TEV at 2% mass ratio of expected yield (e.g. 2mg per 100mg protein). The TEV cleavage was allowed to continue for at least 24 Hrs. The reaction mixture was collected, and several subsequent washes of the resin with TEV buffer were combined with this collection and concentrated for injection onto a size exchange chromatography column (Superdex 200 Increase 10/300 GL, Cytiva) and collected using an ÄKTA pure FPLC (Cytiva), eluting in SEC buffer (30 mM HEPES (pH 7.4), 100 mM NaCl, 10 mM MgCl_2_, 1mM DTT, and 10% glycerol). Fractions containing pure protein were collected and concentrated up to 40mg/mL using Vivaspin Turbo centrifugal concentrators (Sartorius) and snap frozen in liquid nitrogen.

### Detection of covalent modification of LRRK2 RocEXT by whole-protein MS

To quantify labeling of Roc protein by switch II targeting compounds, 2 µM protein was mixed with 50 µM compound in a buffer containing 20mM HEPES pH 7.4, 150mM NaCl, and 1mM MgCl_2_, and immediately subjected to consecutive mass measurements for at least two hours at 20°C via electrospray MS using a Waters Xevo G2-XS system equipped with an Acquity UPLC BEH C4 1.7 µm column. The mobile phase was a linear gradient of 5–95% acetonitrile/water + 0.05% formic acid. Injection time stamps were used to calculate elapsed time. Divarasib (GDC6036) and AMG510 were obtained from MedChemExpress.

### Differential Scanning Fluorometry

DSF experiments were carried out using pre-labeled protein that was desalted using size exclusion. Protein solutions were added to 96 well white plates and then allowed to incubate with added ligand/nucleotide for 15 minutes on ice. Then, SYPRO solution was added, and plates were quickly sealed and spun down at 200xg for 10 seconds prior to starting the experiment. Final concentrations were 4 µM protein, and 4X SYPRO Orange (Invitrogen). Readouts were obtained using a Biorad C1000 Touch thermocycler coupled to a CFX96 RT-PCR system, with a ramp rate of 1C°/min from 4°C to 70°C. Raw values obtained from these experiments were normalized and then analyzed using either Boltzmann sigmoidal function in Graphpad Prism or with dsfworld.com. Experiments were designed quadruplicate, with each condition matched to a same-plate non-protein control. No baseline subtraction was performed.

### Cellular LRRK2 Expression and drug treatment

LRRK2 cDNA was inserted into a pSF-CMV-NEO-NH2-3XFLAG (Sigma-Aldrich) backbone using Gibson assembly cloning. HEK293T cells were obtained from ATCC, propagated, and seeded at 4-5x10^5^ cells per well in six-well plates coated with poly-D-Lysine (Corning). After 24 hours, transient transfections of 1-2 ug DNA/well were carried out using Fugene 6 (Promega), following manufacturer directions at a ratio of 1µg DNA: 2µL Fugene 6. After 24 hours, media was replaced with media containing drugs if necessary and incubated for 8 hours prior to lysis.

### Western Blot Analysis

Western blots were performed as previously described: (dx.doi.org/10.17504/protocols.io.ewov14znkvr2/v2). Briefly, HEK293T cells were subjected to treatments then chilled on ice and washed with ice-cold PBS. Pre-chilled lysis buffer (50 mM Tris-HCl pH 7.5, 270 mM sucrose, 1mM EGTA, 1mM EDTA, 1% Triton X-100) prepared with 1 mM microcystin-LR, 2 mM PMSF, and 1.5X HALT protease and phosphatase inhibitor was added to cells and allowed to incubate for 10 minutes on ice. Cells were scraped and collected into pre-chilled microtubes, vortexed every 5 minutes for 15 minutes, then centrifuged at 20,000xg for 15 minutes at 4◦C. The resultant supernatant was isolated and protein concentration was determined using Pierce BCA assay. 30 µg of protein per sample was loaded into a 1.5 mm 4-12% bis-tris acrylamide gradient gel (Invitrogen) and SDS-PAGE was run in MES running buffer (Invitrogen) at 180V for 70 min. Protein bands were transferred to 0.2µM nitrocellulose membranes using dry transfer (Invitrogen iBlot3) at 25 V for 6 minutes. The resultant membrane was blocked in Intercept® TBS Blocking Buffer for 1 hour at 24◦C. Primary antibody binding was performed overnight at 4◦C with the indicated antibodies diluted in Intercept® TBS Antibody Diluent. After washing membranes for 10 minutes three times with TBST, secondary antibodies diluted 1:10,000 in Intercept® TBS Antibody Diluent were added to membranes and incubated at 24◦C for 1 hour. The membranes were washed for 10 minutes three times with TBST and imaged on a BioRad Chemidoc imager.

### Protein Crystallization

RocEXT-DS bound by GDP and purified by size exclusion chromatography was diluted to 20 mg/mL in SEC buffer (see above). Divarasib was added as a 50 mM solution in DMSO to a final concentration of 1mM. The mixture was placed in a tube rotator at 4 °C for 12 hours, or until LC-MS analysis of the reaction mixture confirmed complete conversion to a single covalent adduct. The reaction mixture was spun down at 20,000 x g for 5 minutes to remove any precipitants. The resultant supernatant was subjected to size exclusion chromatography and concentrated to 40 mg/mL. 0.1 µL of the purified protein-divarasib adduct was mixed with 0.1 µL of well buffer containing Morpheus Fusion (Molecular Dimensions) condition C7, consisting of 30 mM Halides (10mM each of NaF, NaBr, and NaI), 0.12 M Monosaccharide mix 1 (0.02 M each of D-Glucose, D-Mannose, D-Galactose, L-Fucose, D-Xylose, and N-Acetyl-D-Glucosamine), 0.1 M Buffer System 1 pH 6.5 (MES + imidazole buffer), and 30% Precipitant mix 1 (20% PEG 500 MME, 10% PEG 20,000). Drops were set up using the sitting drop method at 4°C, with or without seed crystals. Maximal crystal growth was observed within 2 weeks. The crystals were mounted on nylon loops and flash frozen in liquid nitrogen without additional cryoprotectant solution.

### X-ray data collection and structure determination

The dataset was collected at 100 K at the Advanced Light Source (beamline 8.2.1) at a wavelength of 1.02808 Å using a Dectris EIGER2 Si 9M detector. The dataset was indexed and integrated using iMosflm, scaled with Aimless (CCP4 suite). The structure was solved by molecular replacement using Phaser-MR in Phenix software suite, using the crystal structure of GDP-bound LRRK2 RocEXT (PDB: 6OJE) as the search model. Ligand restraints and coordinates were generated using JLigand. Iterative rounds of manual model rebuilding and reciprocal space refinement were performed using Coot and phenix.refine (v1.20.1-4487) The final model was refined at 2.3 Å resolution with Rwork of 22.55% and an Rfree of 26.15% (99.62% overall completeness). Stereochemical validation via MolProbity demonstrated high model quality with 98.48% of residues in Ramachandran favored regions, 1.52% in allowed regions, and 0.00% outlieers. Complete data collection and refinement statistics are summarized in Table 1. Atomic coordinates and structure factors have been deposited in RCSB Protein Data Bank (PDB: 9C76).

## Author Contributions

R.P., L.Z., K.Z.G., and K.M.S. conceived the project. L.Z. performed protein purification, *in vitro* characterization, and cryo-EM work (data collection and analysis). R.P. performed the cell culture assays and analyzed the data. V.B., H.W., and J.M. performed experiments related to this study and contributed to data interpretation. R.P., L.Z. and K.M.S. wrote the manuscript with input from all authors.

## Acknowledgements

We would like to thank Andres Leschziner and Samara Reck-Peterson for helpful discussions related to LRRK2 conformational control and Dario Alessi, Jia Jun, and Francesca Tonelli for helpful discussions about LRRK2 downstream readouts as well as ongoing advice on generation of cellular models of LRRK2 variants. We would also like to thank the staff at the Advanced Light Source beamline 8.2.1 for help with cryo-EM data collection and processing. L.Z. was supported by a F31 Fellowship from the National Institutes of Health (F31NS122434). V.B. is supported by a F31 Fellowship from the National Institutes of Health (F31CA301949). The study is funded by the joint efforts of the Michael J Fox Foundation for Parkinson’s Research (MJFF) and the LRRK2 Investigative Therapeutics Exchange (LITE) initiative. The University of Dundee administered the grant (MJFF-025924) on behalf of LITE and itself.

