## Supplementary Figures for "Chemical Genetic Targeting of the LRRK2 GTPase Domain"

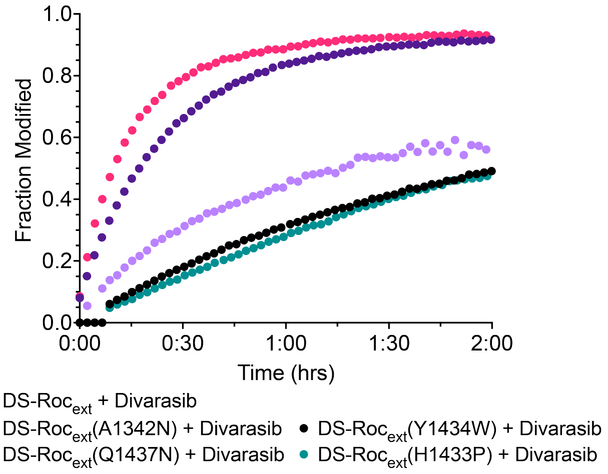

**Supplementary Figure 1:** Reversion of individual drug-sensitizing mutations diminishes divarasib covalent labeling of DS-Roc$_{ext}$.

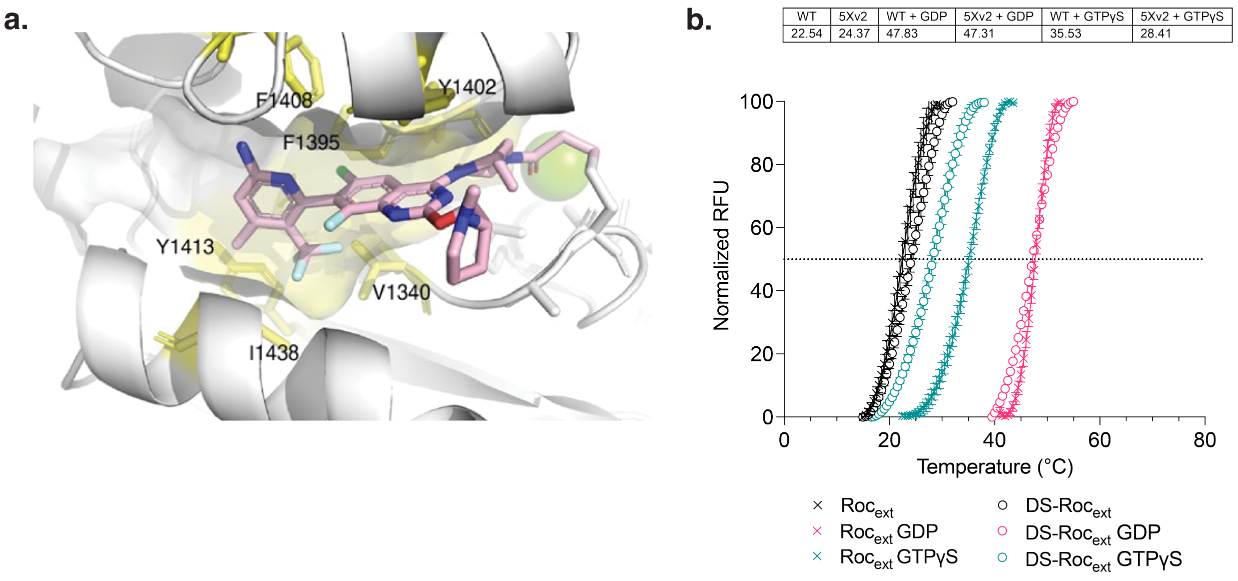

| WT | 5Xv2 | WT + GDP | 5Xv2 + GDP | WT + GTPγS | 5Xv2 + GTPγS |
|---|---|---|---|---|---|
| 22.54 | 24.37 | 47.83 | 47.31 | 35.53 | 28.41 |

**Supplementary Figure 2: A)** CryoEM structure of DS-Roc$_{ext}$ with represented native residues (yellow) contributing to the deep hydrophobic pocket SII pocket. **B)** Comparison of WT-Roc$_{ext}$ and DS-Roc$_{ext}$ DSF performed in nucleotide-saturating conditions.

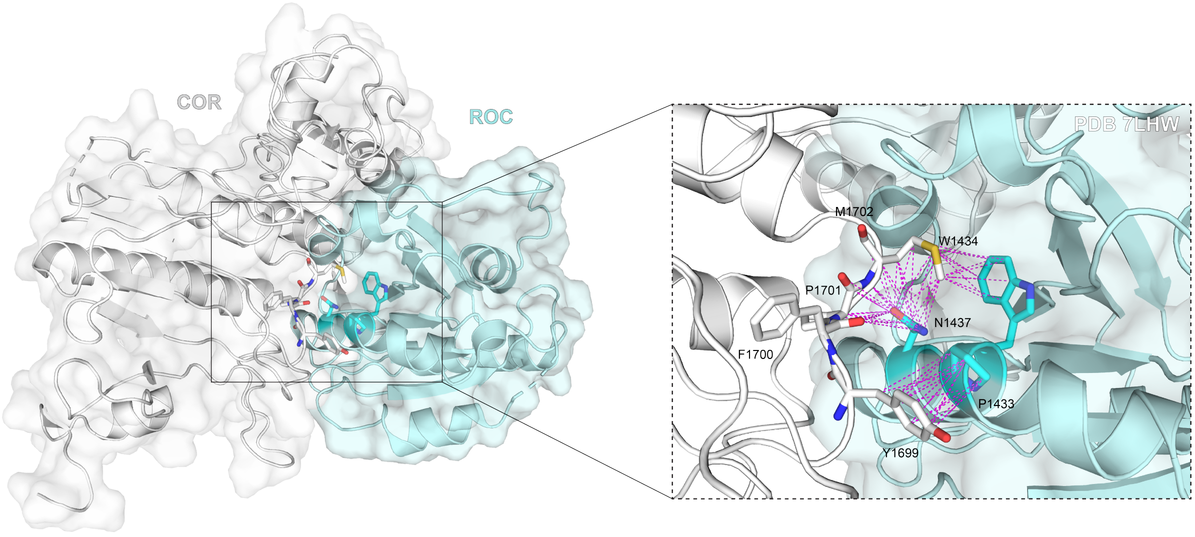

**Supplementary Figure 3**: Drug-sensitizing mutations on the α3 helix contact COR-B residues within a 5Å distance.

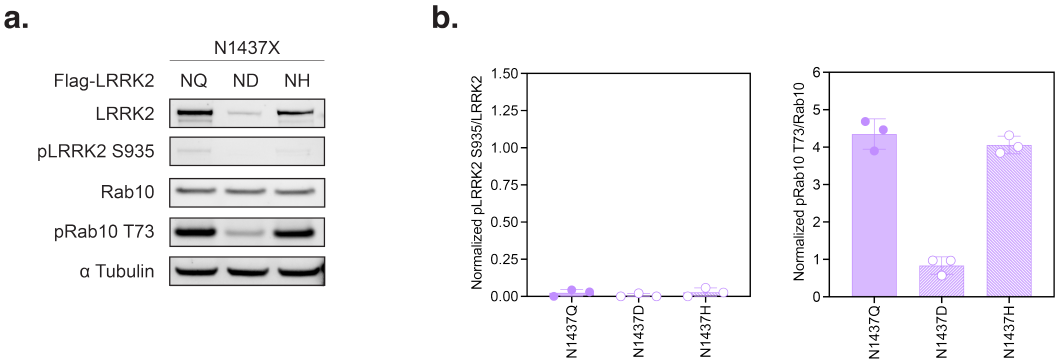

**Supplementary Figure** 4: Drug-sensitizing mutation N1437Q elevates kinase activity in DS-LRRK2. **A)** Immunoblot analysis of HEK293T cells overexpressing FLAG-LRRK2 containing mutation N1437Q (NQ), N1437D (ND), or N1437H (NH). **B)** Immunoblot quantification of LRRK2 S935 phosphorylation and Rab10 T73 phosphorylation for immunoblot shown in Supplementary Figure 4A. Immunoblot quantifications are mean +/- SD (n=3 independent experiments).

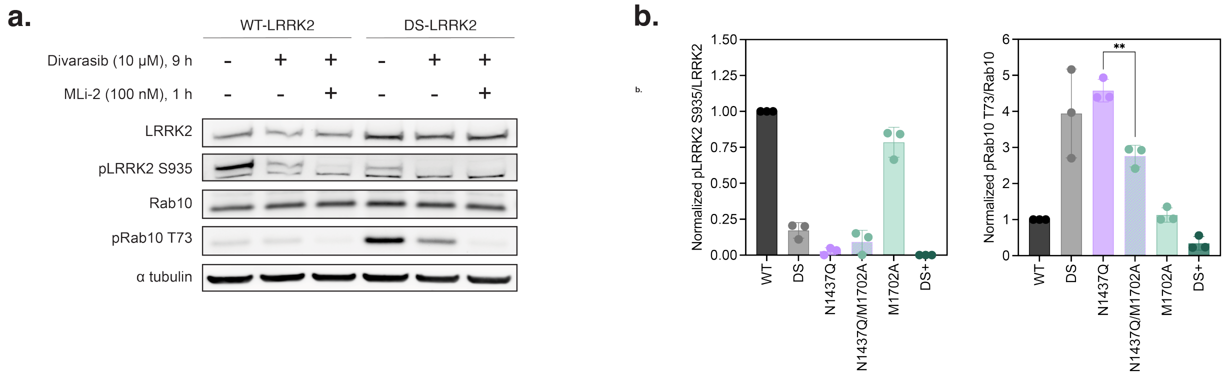

**Supplementary Figure 5**: **A)** Immunoblot analysis of HEK293T cells overexpressing WT-FLAG-LRRK2 or DS-FLAG-LRRK2 sequentially treated with 10 µM Divarasib for 8 hours then 100 nM MLi-2 for 1 hour. **B)** Immunoblot quantification of LRRK2 S935 phosphorylation and Rab10 T73 phosphorylation for immunoblot shown in Figure 4E. Immunoblot quantifications are mean +/- SD (n=3 independent experiments).

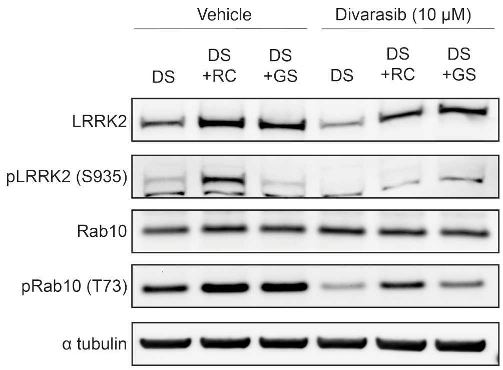

**Supplementary Figure 6.** Immunoblot analysis of HEK293T cells overexpressing DS-FLAG-LRRK2, DS-FLAG-LRRK2 with R1441C (DS+RC), or DS-FLAG-LRRK2 with G2019S (DS+GS) treated with 10 µM divarasib for 8 hours.

**Supplementary Table 1** Crystallographic data collection and refinement statistics.

| Parameter | Value |
| --- | --- |
| **Data Collection** | |
| Space group | $P2_12_12_1$ |
| Unit cell dimensions | |
| $a, b, c$ (Å) | 40.41, 98.38, 105.20 |
| $\alpha, \beta, \gamma$ (°) | 90.00, 90.00, 90.00 |
| Resolution range (Å)* | 49.19–2.10 (2.16–2.10) |
| $R^*_{\mathrm{merge}}$ | 0.319 (2.227) |
| $\mathrm{CC}^*_{1/2}$ | 0.977 (0.375) |
| $\langle I/\sigma(I)\rangle^*$ | 7.9 (1.2) |
| Completeness (%)* | 99.8 (100.0) |
| Redundancy / Multiplicity* | 6.2 (6.4) |
| **Refinement** | |
| Resolution range (Å) | 35.22–2.30 |
| No. reflections (work / test) | 17,332 / 1,935 |
| $R_{\mathrm{work}}/R_{\mathrm{free}}$ | 0.2255 / 0.2616 |
| Wilson $B$-factor | 41.59 |
| Overall mean | 52.32 |
| Protein (Chain A / Chain B) | 61.77 / 44.01 (Mean: 52.36) |
| Ligand (GDP / LRJ) | 58.27 / 53.43 (Chain D: 65.79, Chain G: 41.06) |
| Solvent (Water, 101 atoms) | 46.95 |
| Bond lengths (Å) | 0.002 |
| Bond angles (°) | 0.580 |
| Clashscore | 5.87 |
| **Ramachandran analysis** | |
| Favored (%) | 98.48 |
| Allowed (%) | 1.52 |
| Outliers (%) | 0.00 |
| Rotamer outliers (%) | 1.49 |
| $C_\beta$ deviations (%) | 0.00 |
| Coordinate error (ML-based, Å) | 0.32 |

**Supplementary Table 2** Tm comparison of WT-$Roc_{ext}$ and DS-$Roc_{ext}$ in nucleotide-saturating conditions. Temperatures listed in ˚C.

| | WT-$Roc_{ext}$ | DS-$Roc_{ext}$ |
|---|---|---|
| No nucleotide | 22.54 | 24.37 |
| GDP | 47.83 | 47.31 |
| GTPγS | 35.53 | 28.41 |
